# Multi-tissue transition of A-to-I RNA editing pattern and its regulatory relevance in transcription, splicing, and translation during development

**DOI:** 10.1101/2024.02.21.581478

**Authors:** Jia-Qi Pan, Xu-Bin Pan, Yan-Shan Liu, Yun-Yun Jin, Jian-Huan Chen

## Abstract

Previous studies have shown that A-to-I RNA editing can occur in various organs and tissues under normal physiological conditions. However, the dynamics of RNA editing and its functional relevance in multiple tissues and organs during the embryo-to-adult transition in mammals remain to be elucidated. Here, we performed a comprehensive analysis of RNA-Seq and Ribo-Seq data from six mouse tissues at embryonic and adult stages to elucidate the A-to-I RNA editing transition during development and to validate it in additional mouse brain datasets spanning multiple developmental stages. Our results revealed a general transition of up-regulated A-to-I RNA editing activity across numerous tissue types during embryonic-adult development, indicated by significantly increased average RNA editing levels. Consistently, differential RNA editing (DRE) analysis showed more up-regulated than down-regulated RNA editing sites in all six tissue types. Furthermore, such RNA editing transitions during development could contribute to differential gene expression (DEG) at both transcriptional and translational levels, as well as differential alternative splicing (DAS) across multiple tissues. Differentially edited genes with DEG or DAS could be involved not only in tissue-specific biological functions but also in common routine biological processes during development. Notably, *Adarb1* was found to be a more important player than *Adar* in the CNS (the brain and retina), with its expression levels increasing gradually and correlating with the A-to-I RNA editing activity. Our study demonstrates the potential role of A-to-I editing during development across multiple tissues, providing new insights into its regulatory relevance in both transcriptional and translational landscapes.

## Introduction

RNA editing is a post-transcriptional mechanism that alters the transcript sequence and can increase transcript and protein diversities (Tan et al. 2017b). Adenosine-to-inosine (A-to-I) RNA editing, catalyzed by the adenosine deaminases acting on RNA (ADARs) family, is a canonical type of RNA editing in mammals (Bass 2002; Nishikura 2010). A-to-I RNA editing is involved in diverse biological processes, depending on the editing location. RNA editing profiles may vary among tissues and can also dynamically regulate critical biological processes, such as brain development, cancers, and metabolic disorders (Hwang et al. 2016; Kung et al. 2018; Xu et al. 2018; Buchumenski et al. 2021).

Mammals have almost identical genome sequences but exhibit substantial functional diversity across tissues (Yanai et al. 2006; Ong and Corces 2011; Battle et al. 2017; Sonawane et al. 2017). Correspondingly, A-to-I RNA editing in different tissues and developmental stages may also present complex landscapes and functions (Yao et al. 2019). Therefore, the study of A-to-I RNA editing during mammalian development could not only reveal the dynamic regulation of RNA editing but also help unravel the complex mechanisms of this process. Previous studies have shown that A-to-I editing plays a crucial role in brain and neurological development, particularly in ion channels, neurotransmitter receptors, and synaptic plasticity, and contributes to the complexity and adaptability of the nervous system (Hwang et al. 2016; Yang et al. 2021). Recent genome-wide characterization of RNA editing highlighted roles of editing events of glutamatergic synapse during retinal development (Li et al. 2022). However, most of these studies have focused on RNA editing in a single tissue or organ (Yang et al. 2024), and further investigation is needed to compare RNA editing and its functional relevance across different tissues or organs, including transcription, splicing, and translation.

In the current study, we performed comprehensive transcriptome and translatome analyses of six mouse tissues at both embryonic and adult stages. Our research findings suggest a substantial regulatory role for A-to-I RNA editing in transcription, splicing, and translation, providing new insights into its potential effects on both transcriptional and translational landscapes during development.

## Results

### Tissue-specific A-to-I RNA Editing across mouse development

From the RNA-seq data of six tissues at E15.5 and P42, we identified a total of 618 high-confidence A-to-I RNA editing sites in 317 genes in brain, 501 high-confidence A-to-I RNA editing sites in 265 genes in the heart, 503 high-confidence A-to-I RNA editing sites in 254 genes in the kidney, 479 high-confidence A-to-I RNA editing sites in 235 genes in the liver, 660 high-confidence A-to-I RNA editing sites in 323 genes in the lung, 853 high-confidence A-to-I RNA editing sites in 404 genes in the retina **(Figure 1A; Supplementary Table S1)**. Analysis of the functional categories showed that the majority (68.5-86.4%) of them were 3′-untranslated region (3′UTR) RNA editing. Missense RNA editing accounted for only a small proportion (2.7-7.3%) of the total variants **(Figure 1B, Supplementary Figure S1A)**. Sorting intolerant from tolerant (SIFT) analysis predicted that 15.4-33.3% of missense variants could potentially impact the function of the encoded protein **(Figure 1C, Supplementary Figure S1B)**. Motif analysis showed that G was suppressed at one base upstream but preferred at one base downstream of the editing sites **(Figure 1D)**, consistent with the reported sequence preference of *ADAR* editing.

**Figure 1.**
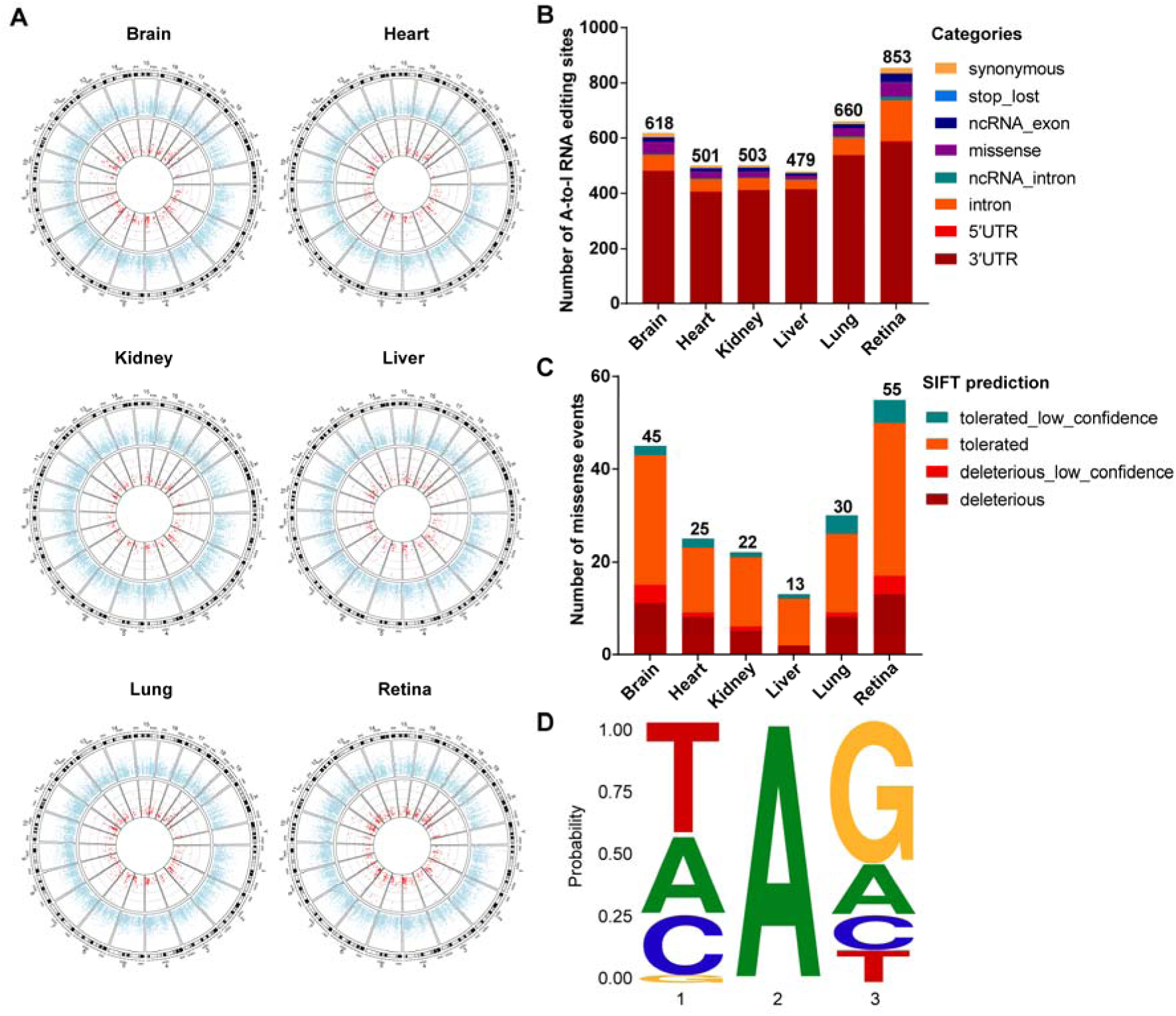
Transcriptome-wide analysis of A-to-I RNA editing in different tissues from embryonic (E15.5) and adult (P42) mice. **(A)** Circos plot of gene expression and A-to-I RNA editing sites in all six tissues. The outer circle denotes the mean expression level of genes, and the inner circle denotes the A-to-I RNA editing sites. **(B)** Functional categories of A-to-I RNA editing variants in all six tissues. **(C)** SIFT prediction of missense A-to-I RNA editing variants. **(D)** Motif context around A-to-I RNA editing sites.

To investigate the dynamic regulation of A-to-I RNA editing, we compared the existence of editing sites and edited genes between E15.5 and P42 in different tissues. As shown in **Figure 2A and Supplementary Figure S2**, more editing sites and edited genes were observed at P42 than at E15.5 across different tissues, except in the heart.

**Figure 2.**
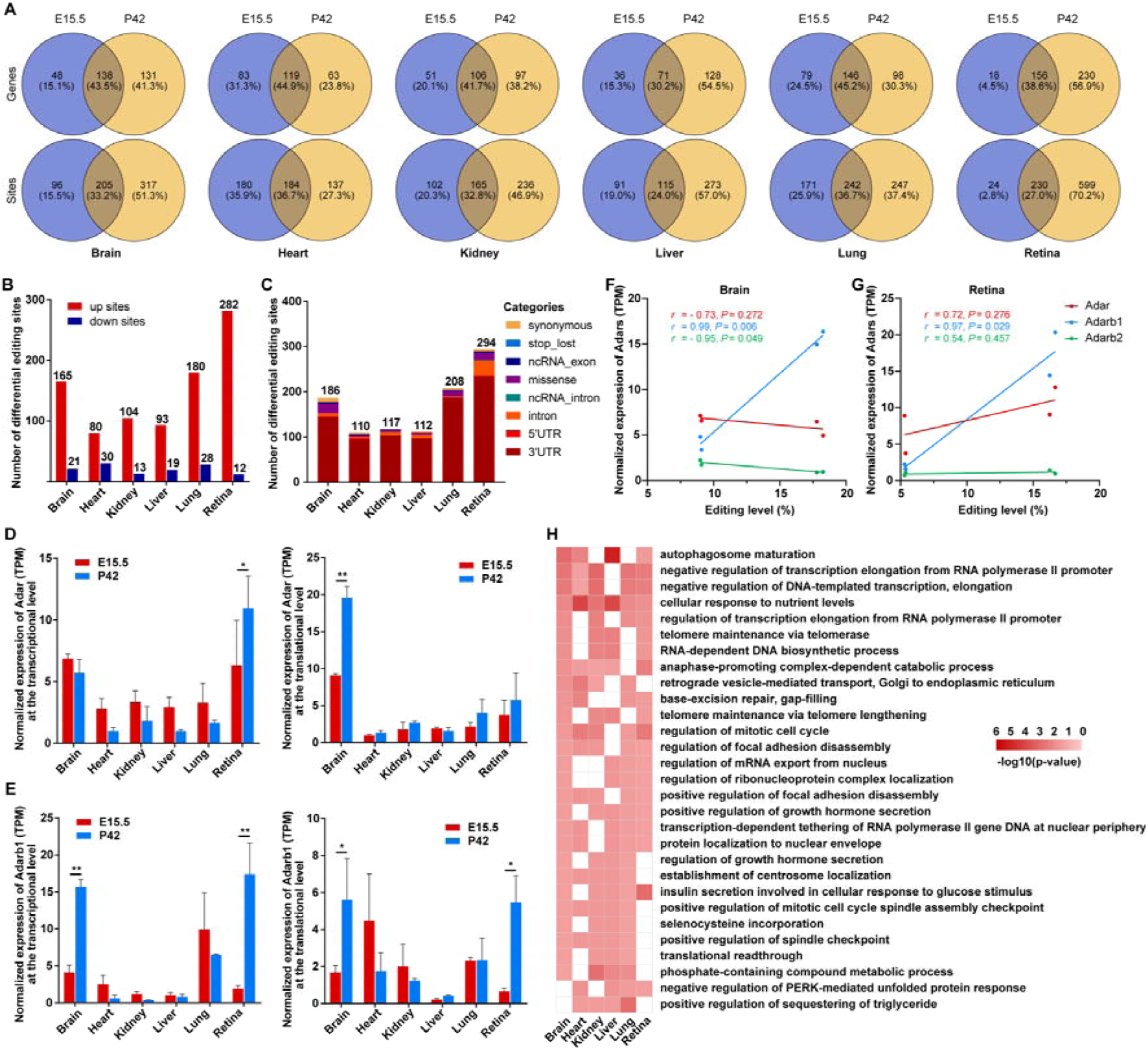
Differential A-to-I RNA editing in different tissues between embryonic (E15.5) and adult (P42) mice. **(A)** Venn plots of genes and sites with A-to-I RNA editing at embryonic and adult stages. **(B)** Bar plot showing the number of DRE sites between embryonic and adult stages. **(C)** Functional categories of DRE sites. **(D)** Normalized expression of *Adar* at the transcriptional (left) and translational level (right). **(E)** Normalized expression of *Adarb1* at the transcriptional (left) and translational level (right). **: *P* < 0.01; *: *P* < 0.05. *Pearson*’s correlation analysis between editing levels and normalized expression of *Adars* at the transcriptional level in the brain **(F)** and the retina **(G)**. **(H)** Heatmap showing terms from GO items enriched by differential edited genes in ≥ three tissues.

### Differential A-to-I RNA Editing between embryonic and adult stages

To systematically analyze RNA editing events during development, we then analyzed differential RNA editing (DRE) between embryonic and adult stages. We identified 186, 110, 117, 112, 208, 294 DRE sites in these six tissues, respectively **(Figure 2B; Supplementary Table S2)**. The top 50 DRE sites across different tissues, ranked by p-values, are shown in **Figure 3**. In addition, we found that most DRE sites were up-regulated at P42 compared with E15.5. Consistent with the overall RNA editing landscape, most of the DRE sites were also found in 3UTR **(Figure 2C)** and showed similar sequence motifs across all tissues **(Supplementary Figure S3A)**.

**Figure 3.**
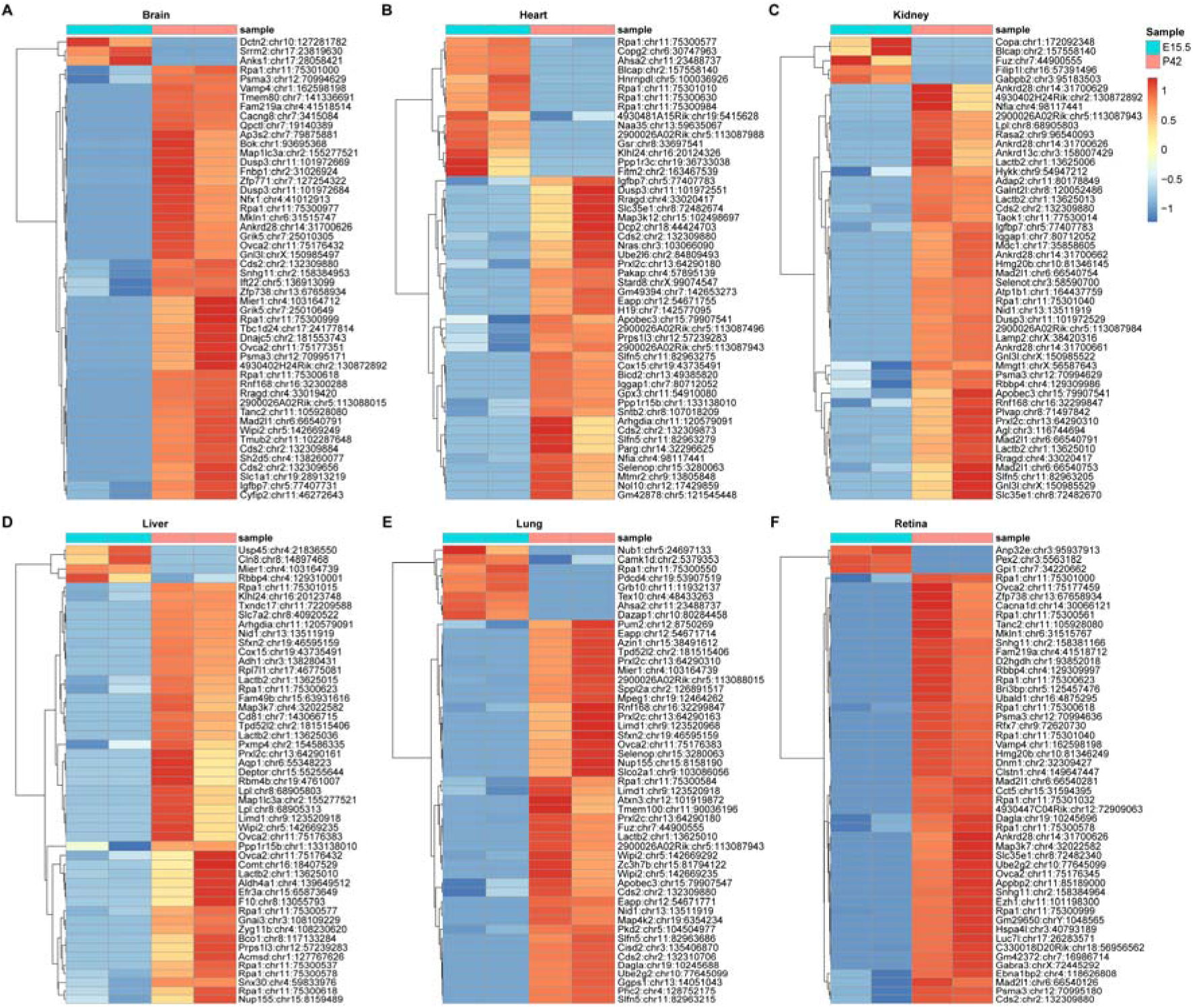
A-to-I DRE sites in different tissues during development. Heatmap showing the top 50 editing levels of DRE sites in different tissues during development.

We then evaluated the expression of Adars across different tissues at both stages. Our findings revealed a dramatic increase in Adarb1 expression during brain and retinal development at both transcriptional and translational levels **(Figure 2E)**. *Adar* was significantly up-regulated in the brain at the translational level but not at the transcriptional level, and in the retina at the transcriptional level but not at the translational level **(Figure 2D)**. In addition, we observed a significant positive correlation between the mean editing level and *Adarb1* mRNA expression both in the brain and retina (*Pearson*’s *r* = 0.99, *P* = 0.006, and *Pearson*’s *r* = 0.97, *P* = 0.029, respectively) **(Figure 2F, G)**. We then identified *Adarb1*-correlated DRE sites by evaluating the correlation of their editing levels with *Adarb1* mRNA expression levels in the brain and retina. 11 *Adarb1*-correlated DRE sites in 11 genes. 18 ones in 13 genes were identified in the brain and retina, respectively, all of which sites were significantly up-regulated and positively correlated with *Adarb1* expression (**Supplementary Figure S4, S5**). These observations indicated that *Adarb1* may be the primary editing enzyme of A-to-I editing sites in the brain and retina. However, we failed to observe a significant correlation between editing levels and *ADAR* expression in the other four tissues **(Supplementary Figure S3B and C)**.

### Functional enrichment analysis of differentially edited genes

To explore the biological function of tissue-specific A-to-I RNA editing in development, we performed enrichment analysis of DRE genes in each tissue. GO analysis indicated that these DRE genes were not only enriched for certain tissue-specific biological functions but also various routine biological processes related to development, such as ‘positive regulation of growth hormone secretion’, ‘regulation of mitotic cell cycle’, ‘RNA-dependent DNA biosynthetic process’, and ‘phosphate-containing compound metabolic process’ **(Figure 2H; Supplementary Table S3)**.

### Potential effects of A-to-I RNA editing on transcription and translation

To explore the landscape function of A-to-I RNA editing at transcriptional and translational levels, we compared the DRE genes and differentially expressed genes (DEGs) identified by mRNA-Seq or Ribo-seq in different tissues **(Figure 4A; Supplementary Table S4)**. Our results revealed that 46, 18, 40, 37, 59, and 58 DRE genes were differentially expressed both in mRNA-Seq and Ribo-Seq in the brain, heart, kidney, liver, lung, and retina, respectively **(Figure 4B)**. We then evaluated these DRE genes with differential expression for significant functional enrichment in at least two tissue types. DRE genes with up-regulated expression were enriched in routine biological processes of protein modification and metabolic regulation, such as ‘protein-containing complex localization’, ‘dephosphorylation’, ‘regulation of lipid metabolic process’, and ‘glycerolipid metabolic process’. In contrast, DRE genes with down-regulated expression were enriched in biological processes related to DNA and RNA dynamics, such as ‘DNA conformation change’, ‘DNA replication’, ‘mRNA export from nucleus’, and ‘RNA localization’ **(Figures 4C, D; Supplementary Table S5)**.

**Figure 4.**
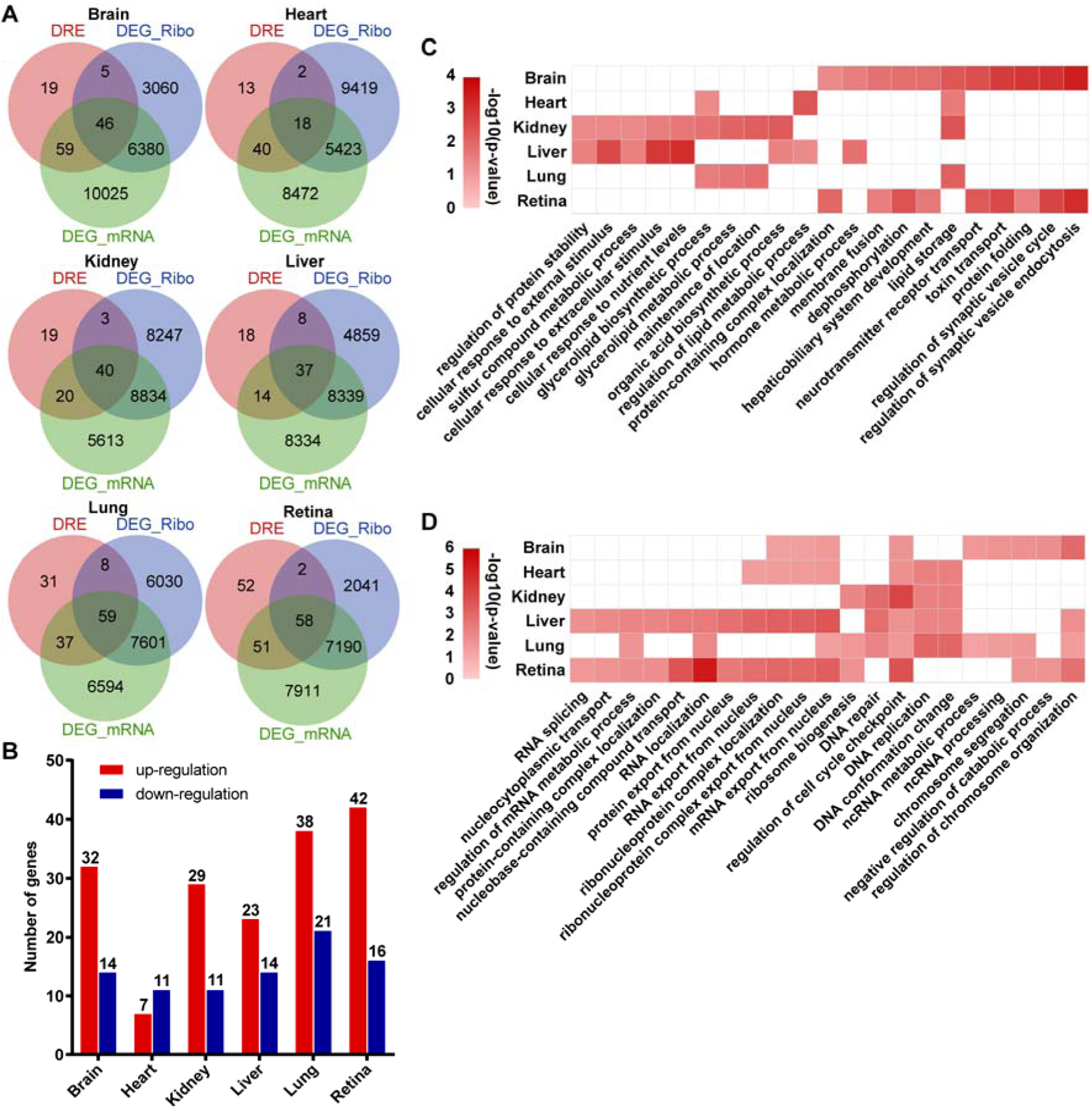
Possible functional impact of A-to-I DRE on transcription and translation during development. **(A)** Venn plots comparing genes with DRE and DEGs identified by mRNA-seq (DEG_mRNA) or Ribo-seq (DEG_Ribo) across different tissues. **(B)** Bar plot showing the number of DRE genes differentially expressed both in mRNA-Seq and Ribo-Seq. Heatmap showing the enriched GO terms for DRE genes with up-regulated **(C)** and down-regulated expression **(D)** in at least two tissue types.

In addition, it was also noticed that 59, 40, 20, 14, 37, and 51 DRE genes were differentially expressed at the transcription level only but not at the translation level in the brain, heart, kidney, liver, lung, and retina, respectively **(Supplementary Figure S6A)**. We then evaluated these DRE genes with differential RNA expression for significant functional enrichment in at least two tissue types. Likewise, our enrichment analysis indicated that these DRE genes, with transcriptional differential expression only, were enriched for biological processes involved in the regulation of cellular processes and metabolic processes, such as ‘cellular component disassembly’, ‘negative regulation of cell motility’, ‘thioester metabolic process’, and ‘nucleoside bisphosphate metabolic process’. In contrast, these DRE genes with down-regulated expression were mainly enriched for biological processes involved in DNA regulation and gene silencing, such as ‘positive regulation of DNA repair’, ‘positive regulation of DNA metabolic process’, ‘regulation of gene silencing’, and ‘positive regulation of posttranscriptional gene silencing’ **(Supplementary Figure S6B, C; Supplementary Table S4)**.

### RNA binding prediction of differential A-to-I RNA editing sites

To gain insights into the potential functional impact of RNA editing, we utilized the RBPmap database to predict the binding patterns of RBP overlapping with differential A-to-I RNA editing sites across different tissues **(Supplementary Figure S7)**. **Figure 5** illustrates the top 10 RBPs with the highest binding frequencies to differential A-to-I RNA editing sites across various tissues. Notably, RNA binding motif protein 45 (*RBM45*), Heterogeneous nuclear ribonucleoprotein A0 (*HNRNPA0*), Serine and arginine-rich splicing factor 10 (*SRSF10*), Unk Zinc Finger (*UNK*), and CCR4-NOT transcription complex subunit 4 (*CNOT4*) were identified across all six tissues. This suggests that these RBPs may play a crucial and ubiquitous role in regulating RNA editing, indicating potential functional significance across gene expression and cellular processes.

**Figure 5.**
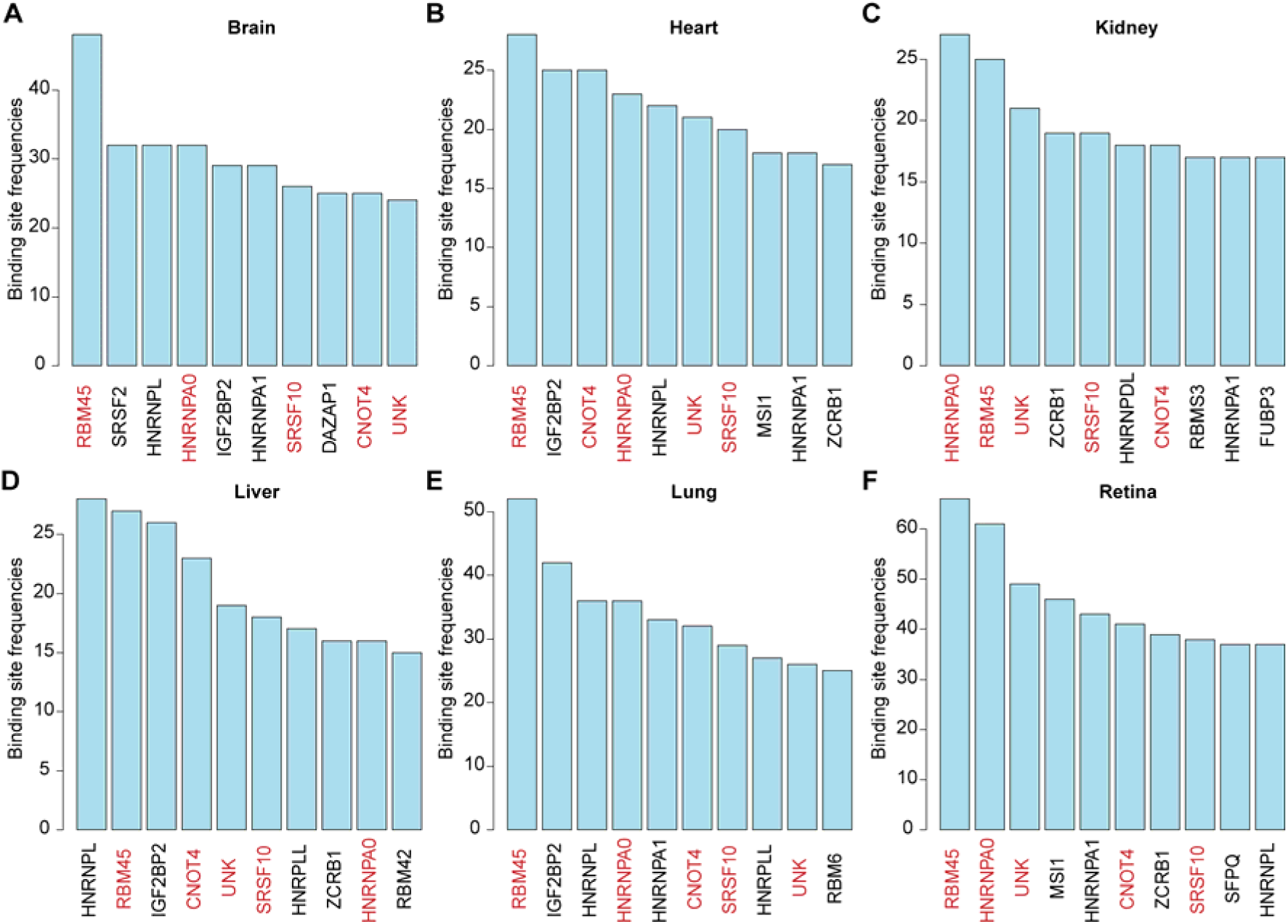
The top 10 frequent RNA binding proteins (RBPs) ranked by their binding frequencies to DRE sites in different tissues.

### Interaction between A-to-I RNA editing and alternative splicing

To gain insights into the interplay between A-to-I RNA editing and alternative splicing, we used rMATS to identify alternative splicing (AS) and differential alternative splicing (DAS) events across tissues. We found that 39-54% of genes with A-to-I RNA editing also exhibited alternative splicing in these six tissues, suggesting a potential interaction between A-to-I RNA editing and alternative splicing **(Figure 6A)**. Then, we identified DAS events and genes that differed between the embryonic and adult stages **(Figure 6B; Supplementary Table S6)**. Among the six tissues, the brain and retina exhibit the highest numbers of DAS events and genes. Comparing DRE genes and DAS genes, we found that 4-40 DAS genes were also differentially edited in these six tissues during development **(Figure 6C)**. Enrichment analysis indicated that these genes were involved in various routine biological processes related to development, such as ‘negative regulation of programmed necrotic cell death’, ‘positive regulation of cytoplasmic transport’, ‘fibroblast growth factor receptor signaling pathway’, and ‘neuron cell-cell adhesion.’ This suggests that RNA editing and alternative splicing were closely intertwined in cellular processes such as survival, death, signal transduction, and cell-cell interactions during development **(Figure 6D)**.

**Figure 6.**
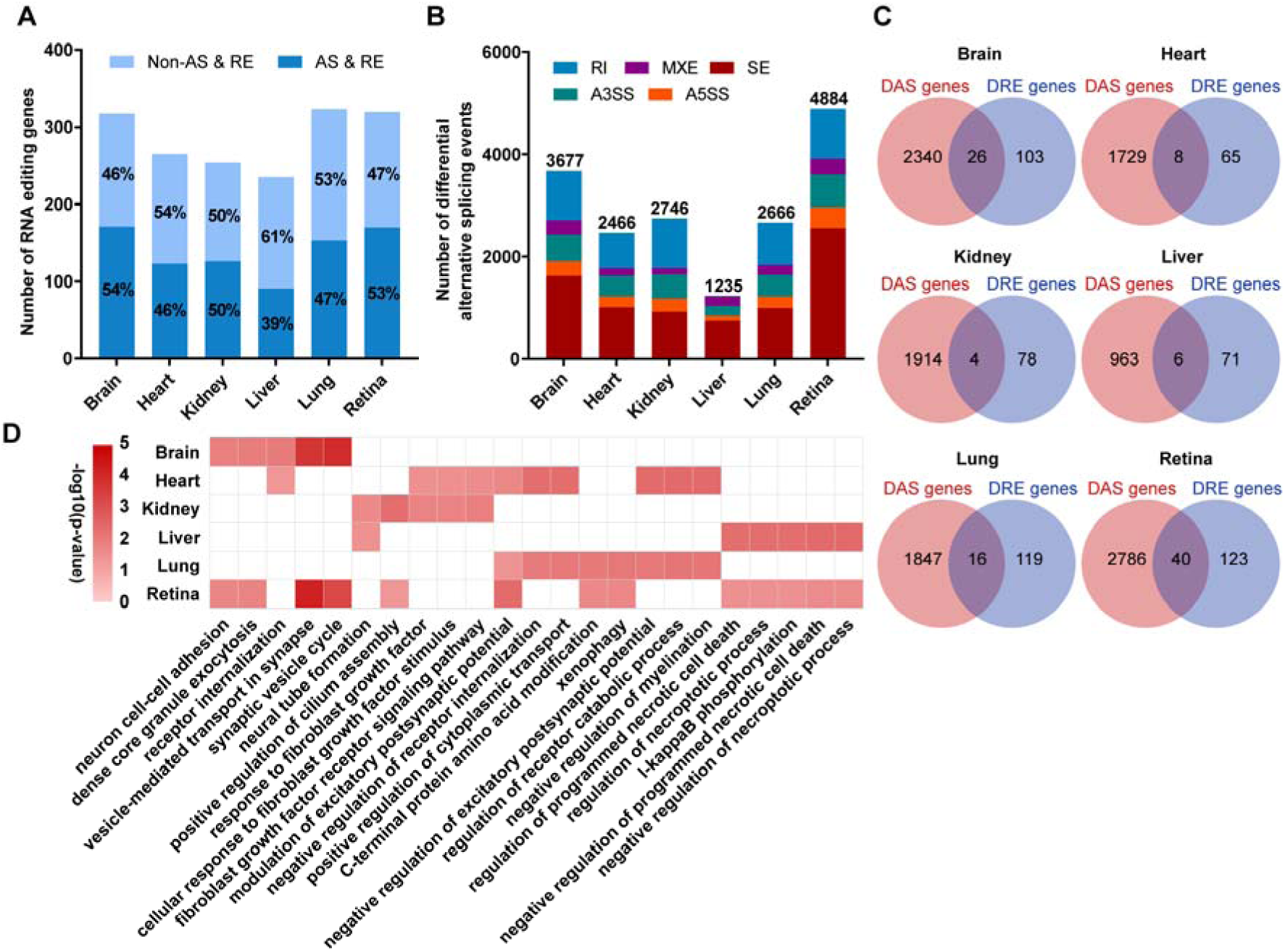
Developmental interrelation between A-to-I RNA editing and alternative splicing. **(A)** Proportion and number of edited genes with and without alternative splicing (AS) in different tissues. **(B)** Bar plot comparing the number of differential alternative splicing (DAS) events between embryonic and adult stages. **(C)** Venn plots of DAS genes and DRE genes in different tissues. **(D)** Heatmap showing the enriched GO terms for DAS genes with differential editing in at least two tissue types. RI, retained intron; MXE, mutually exclusive exons; A3SS, alternative 3′ splice sites; A5SS, alternative 5′ splice sites; SE, skipped exon.

### Validation of A-to-I RNA editing pattern during brain development

To validate the common characteristics of A-to-I RNA editing pattern during development, we conducted a comparative analysis using additional RNA-Seq and Ribo-Seq datasets (GSE157425 and GSE169457) of brain neocortex at embryonic (E12.5–E17) and postnatal (P0) stages from an independent study. Using the same analysis procedure, 557 to 723 A-to-I RNA editing sites were identified across different developmental stages, with a significant increase in the number of editing sites over time. Consistent with the findings in the brain RNA editing data described above, RNA editing was also enriched in 3UTR **(Figure 7A)**. A large proportion (283, 30.01%) of the editing sites were shared across all stages, with a considerable number also shared by adjacent time points, whereas only a small proportion (9.23%, 87) were exclusively found in a single stage **(Figure 7B)**. Analysis of editing levels between different stages revealed an increasing trend of the average editing level during brain development **(Figure 7C)**. To systematically explore the temporal dynamics of RNA editing, DRE across these time points was analyzed. Our results showed that 453 sites underwent differential editing during brain development **(Figure 7D and Supplementary Figure S8A)**. The top 50 DRE sites over time, ranked by p-values, are shown in **Figure 7E**. Most DRE sites were up-regulated during development, with the most rapid up-regulation occurring after birth. By comparing the discovery and validation datasets, 52 differentially edited genes and 42 DRE sites were shared between the two **(Supplementary Figure S8B, C)**. Gene function enrichment analysis indicated that the DRE in the validation dataset was mainly involved in chromatin and gene expression regulation, such as ‘negative regulation of organelle organization’, ‘covalent chromatin modification’, and ‘regulation of chromosome organization’ **(Supplementary Figure S8D)**.

**Figure 7.**
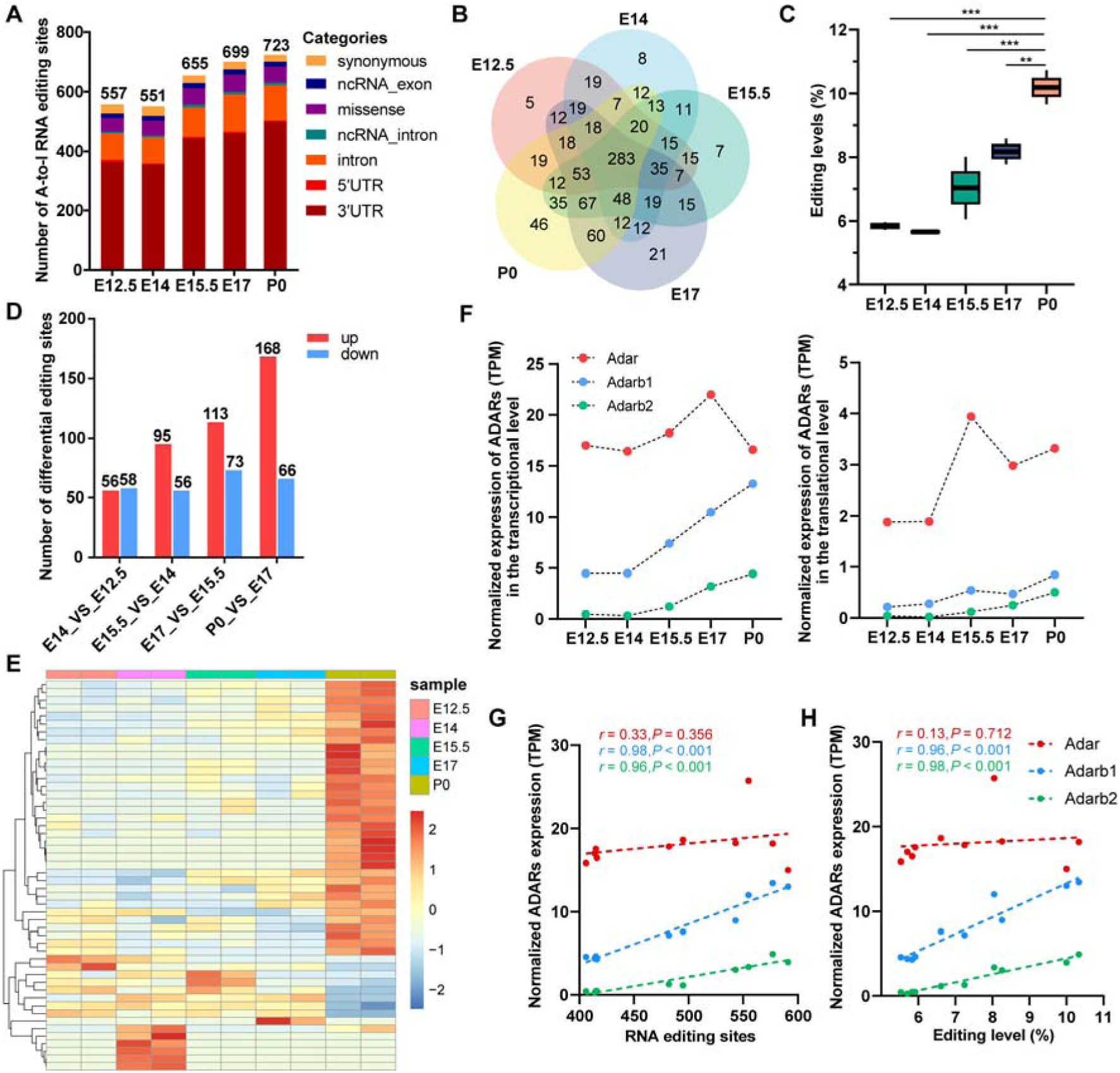
Validation of RNA editing pattern during brain development in dataset GSE157425 and GSE169457. **(A)** Functional categories of A-to-I RNA editing variants in the brain’s neocortex at embryonic (E12.5–E17) and postnatal (P0) stages. **(B)** Intersection of RNA editing sites in each stage. **(C)** Editing levels in each stage. ***: *P* < 0.001; **: *P* < 0.01. **(D)** Bar plot showing the number of differential editing sites between adjacent developmental time points. **(E)** Heatmap showing the top 50 editing levels of DRE sites during brain development. **(F)** Normalized expression of three *Adars* at the transcriptional (left) and translational level (right). **(G)** *Pearson*’s correlation analysis between number of editing sites and normalized expression of *Adars* at the transcriptional level. **(H)** *Pearson*’s correlation analysis between editing levels and normalized expression of *Adars* at the transcriptional level.

Comparison of *Adarb1* and Adarb2 expression among different stages showed a dramatic increase in *Adarb1* and *Adarb2* expression during brain development at both transcriptional and translational levels **(Figure 7F)**. In addition, we observed significant positive correlations of *Adarb1* and *Adarb2* expressions with the number of RNA editing sites and edited genes, as well as with the average editing level (all *Pearson r* range from 0.96 to 0.98 and all *P* < 0.001) **(Figure 7G, H)**.

Then the potential effects of A-to-I RNA editing on transcription and translation were further validated during brain development. We observed that 134 differentially edited genes were differentially expressed both in RNA-Seq and Ribo-Seq over time **(Figure 8A)**. Gene enrichment analysis indicated that differentially edited genes with up-regulated expression were primarily involved in functions related to synapses and neurotransmitters, such as ‘regulation of postsynaptic membrane potential’, ‘receptor diffusion trapping’, and ‘synaptic vesicle cycle’. In contrast, those with down-regulated expression were mainly involved in biological processes related to RNA dynamics, such as ‘positive regulation of mRNA processing’ and ‘regulation of transcription of nucleolar large rRNA by RNA polymerase I’ **(Figure 8B)**. Comparing splicing events between adjacent time points, we revealed 1622∼2741 DAS events over time **(Figure 8D)**. We then found that 85 genes with both DAS and differentially edited during brain development **(Figure 8C),** which were mainly enriched in functions related to synaptic regulation and brain development, such as ‘regulation of neuronal synaptic plasticity’, ‘excitatory postsynaptic potential’, and ‘forebrain development’ **(Figure 8E)**.

**Figure 8.**
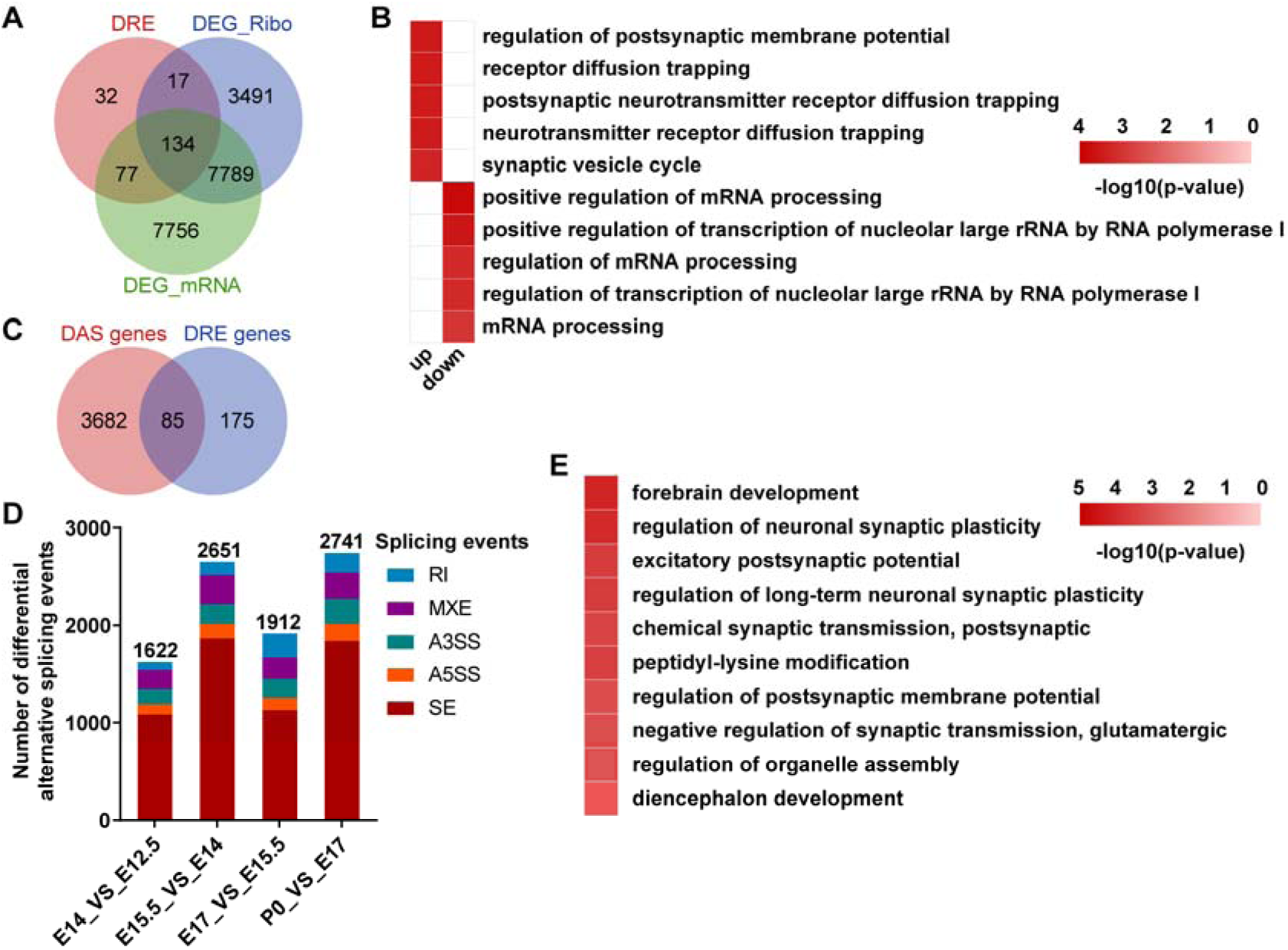
Potential functional impact of A-to-I DRE in transcription, translation, and splicing during brain development. **(A)** Venn plots comparing genes with DRE and DEGs identified by mRNA-seq (DEG_mRNA) or Ribo-seq (DEG_Ribo). **(B)** Heatmap showing the enriched GO terms for DRE genes with up-regulated and down-regulated expression. **(C)** Venn plots comparing DAS genes and DRE genes. **(D)** Bar plot showing the number of DAS events between adjacent developmental time points. **(E)** Heatmap showing the enriched GO terms for DAS genes with differential editing. RI, retained intron; MXE, mutually exclusive exons; A3SS, alternative 3′ splice sites; A5SS, alternative 5′ splice sites; SE, skipped exon.

## Discussion

*ADAR*-mediated A-to-I RNA editing could be crucial for the mammalian development of various organs and tissues (Slotkin and Nishikura 2013). Yet, its regulatory relevance remains to be further elucidated. In the current study, our analysis revealed a dramatic RNA editing transition characterized by increased RNA editing activity across different tissues during embryonic-to-adult development, with potential biological impacts on RNA splicing, transcription, and translation.

Differential missense RNA editing was observed in multiple tissues during the development from E15.5 to P42. Notably, significant up-regulation of two missense editing c.284A>G (p.K95R) and c.232A>G (p.R78G) in the insulin-like growth factor binding protein 7 (*Igfbp7*) gene was observed in the mouse brain, heart, and kidney at the adult stage (P42) compared to the embryonic (E15.5) stage. These two conserved missense editing sites were also found in other mammals, such as humans and pigs (Levitsky et al. 2023). *Igfbp7* encodes a soluble protein that binds insulin growth factor (IGF) and suppresses downstream signaling, thereby inhibiting protein synthesis, cell growth, and survival (Levitsky et al. 2023). Recent studies have reported that K95R and R78G editing in *IGFBP7* are elevated in the adult brain and cardiovascular tissues, as well as in atherosclerosis and cardiomyopathies (Huang et al. 2021; Mann et al. 2023). R/G substitutions in the IGF binding domain may disrupt the interaction between the protein and its IGF ligands (Levanon et al. 2005). In addition, the R78G and K95R recordings have been reported to possibly modulate its susceptibility to proteolysis (Godfried Sie et al. 2012). Our results also showed that another differential missense editing c.14A>G (p.Q5R) in the bladder cancer-associated protein (*Blcap*) gene was observed in the mouse heart, kidney, and retina. The editing level of this site decreased in the heart and kidney, yet increased in the retina, during development from embryo to adult. *Blcap* encodes a tumor suppressor that prevents tumorigenesis by promoting apoptosis and inhibiting cell proliferation. RNA over-editing of *BLCAP* can activate the Akt/mTOR signaling pathway by regulating phosphorylation levels, thereby promoting hepatocellular carcinoma progression (Hu et al. 2015) and cervical carcinogenesis (Chen et al. 2017). In addition, missense editing in Neuroligin 2 (*Nlgn2*) c.406A>G (p.N136D) was up-regulated in the brain, lung, and retina during the embryo-to-adult development, which probably abolishes an N-glycosylation target site in *Nlgn2* (Scott and Panin 2014) and thus might affect neurotransmission. *Nlgn2* encodes a transmembrane scaffolding protein and promotes synapse development by forming trans-synaptic bridges spanning the synaptic cleft (Ali et al. 2020). Moreover, *Nlgn2* has been reported to be associated with neurological and psychiatric disorders, such as schizophrenia (Sun et al. 2011), autism (Parente et al. 2017), and anxiety (Babaev et al. 2016). The functional relevance of most of these missense DRE sites in development, especially those in *Igfbp7* and *Nlgn2,* remains poorly understood. Our findings thus warranted functional investigations in future studies.

Interestingly, our results showed that RNA editing was up-regulated in a subset of long intergenic non-coding RNAs (lincRNAs), such as small nucleolar RNA host gene 11 (*Snhg11*), H19 imprinted maternally expressed transcript (*H19*), and maternally expressed gene 3 (*Meg3*), during development. One intronic editing variant in *Snhg11* and two exonic editing variants in *Meg3* were significantly up-regulated in the brain in adult mice. Seven intronic editing variants in *Snhg11* and four exonic editing variants in *Meg3* were significantly up-regulated in the retina in adult mice **(Supplementary Table S2)**. *Snhg11* has been reported to promote cell proliferation, migration, apoptosis, and autophagy during cancer development (Liu et al. 2020; Huang et al. 2020). Recent studies implicated that *Snhg11* also plays a neuroprotective role in neuronal injury (Chen et al. 2021b). Previous studies have suggested *that Meg3* not only contributes to cancer development but is also associated with the pathological stages of cancer and the prognosis of cancer patients (He et al. 2017). Recent studies have shown that *Meg3* regulates surface AMPA receptor expression in primary cortical neurons and plays a vital role in the nervous system (Tan et al. 2017a). The hyper-editing in *Snhg11* and *Meg3,* and the differential RNA editing in both genes, suggest a potential role for these two lincRNA genes during brain and retina development.

In mammals, A-to-I editing is catalyzed by *ADAR*s, including *ADAR1*, *ADARB1,* and *ADARB2* (Fritzell et al. 2018). Our findings revealed that *Adarb1* was primarily responsible for the dynamics of A-to-I editing during brain and retinal development, compared to *Adar* and *Adarb2*. In particular, *Adarb1* increased gradually and showed a strong correlation with the up-regulated A-to-I RNA editing activity during development, which was further validated in multi-stage brain development datasets. In contrast, the expression of *Adar*, *Adarb1*, and *Adarb2* did not differ significantly across the other four tissues. These results suggest a tissue-specific role for the *ADAR* family in mediating A-to-I RNA editing, highlighting the vital role of *Adarb1* in regulating A-to-I RNA editing activity during CNS development.

Dynamic regulation of transcription and translation plays a key role in embryonic development and maintenance of adult tissues (Matkovich et al. 2014; Falcon et al. 2022). This regulation involves multiple dimensions, with RNA modification standing out as a crucial facet in this intricate process (Roundtree et al. 2017; Cui et al. 2022). Previous studies have shown that A-to-I editing could play an essential role in regulating transcriptome diversity (Bahn et al. 2012). However, the impact of RNA editing on translation during mammalian development remains largely unexplored. Ribo-seq can reveal the intricacies of translation dynamics, enabling exploration of the specific regions of mRNA that are actively undergoing translation by ribosomes (Ingolia et al. 2019). Our results revealed differentially edited genes with differential expression at both mRNA and protein levels, primarily involved in both tissue-specific and routine biological processes related to development. Previous studies have shown that A-to-I editing is crucial in neurological development (Yang et al. 2021). Ectoderm-derived brain and retina are essential components of the CNS (Fernando et al. 2022). Our results consistently showed that DRE may contribute to brain and retinal development by regulating the expression of genes involved in synaptic vesicle and neurotransmitter function, highlighting its crucial role in CNS development.

Previous studies have shown that A-to-I editing of pre-mRNA before RNA splicing could be closely related to alternative pre-mRNA splicing (Hu et al. 2022; Hsiao et al. 2018). However, a global understanding of the interplay between RNA editing and alternative splicing during development remains to be further elucidated. Our study systematically compared differential A-to-I RNA editing genes and differential alternative splicing genes across six tissues, revealing their close intertwining in cell survival, death, signal transduction, and cell-cell interactions during development, providing new insights into the relationship between RNA editing and alternative splicing.

The overlap between differentially edited RNA sites and RBP binding sites provides a potential link between RNA editing and RBP functions in development, such as the regulation of pre-mRNA splicing and higher-order RNA structures (Hu et al. 2022). Our study identified five core RBPs with the highest binding frequencies to DRE sites across different tissues, including *RBM45*, *HNRNPA0*, *SRSF10*, *UNK*, and *CNOT4*. *RBM45* encodes an RBP involved in neural development, and its pathological aggregation is closely related to neurodegenerative diseases, such as amyotrophic lateral sclerosis (ALS) and frontotemporal lobar dementia (FTLD) (Chen et al. 2021a). *SRSF10* encodes an atypical member of serine/arginine-rich (SR) proteins (Zhou et al. 2014), and has been reported to influence glucose, fat, and cholesterol metabolism, embryonic heart development, and neural processes by controlling alternative splicing of specific transcripts (Shkreta et al. 2021). *CNOT4* encodes an evolutionarily conserved E3 ubiquitin ligase (Chalabi Hagkarim and Grand 2020). *CNOT4* has been demonstrated to play synergistic biochemical functions during development and is required for post-implantation embryo development and meiosis progression during spermatogenesis (Dai et al. 2021). Our study suggested that these RBPs are involved in the downstream effects of differential RNA editing during development. Further experimental and functional studies will be needed to further understand the interplay between these RBPs and RNA editing during development.

In conclusion, our study suggested a transition in the A-to-I RNA editing pattern across multiple tissues and its regulatory relevance for transcription, splicing, and translation during development. Further investigation will be needed to evaluate the role and mechanisms of RNA editing within both transcriptional and translational landscapes.

## Materials and methods

### Dataset download

Reads of RNA sequencing (RNA-seq) and ribosome profiling (Ribo-seq) from a dataset (GSE94982) were downloaded from the NCBI Gene Expression Omnibus (GEO, http://www.ncbi.nlm.nih.gov/geo) database. The dataset contains six types of tissue samples from wild-type C57BL/6 mice at embryonic day (E) 15.5 and postnatal day (P) 42 (N = 2 each tissue group per time point), including ectoderm-derived brain and retina, mesoderm-derived heart and kidney, as well as endoderm-derived liver and lung (Wang et al. 2021). The validation datasets (GSE157425 and GSE169457) were downloaded from the NCBI Gene Expression Omnibus (GEO, http://www.ncbi.nlm.nih.gov/geo). The dataset contains neocortex tissue from wild-type CD-1 mice at embryonic (E12.5–E17) and early postnatal (P0) period (N = 2 per time point) (Harnett et al. 2022). The tissue collection, RNA extraction, and library construction and sequencing were performed as described in the original studies.

### Data processing

The raw RNA-Seq and Ribo-Seq sequencing reads were processed using FASTP (version 0.23.4) to remove adapters and low-quality reads (Chen et al. 2018). The obtained Clean data were mapped to the mouse reference genome (UCSC mm10) using RNA STAR (Version 2.7.0e) (Dobin and Gingeras 2015), generating BAM alignment files. SAMtools (Version 1.9) was used to eliminate multiple mappings and optical duplications of reads, retaining reads uniquely mapped to the mouse genome (Li et al. 2009). GATK (Version 4.1.3) was used to recalibrate the base quality scores of the BAM files by following the best practices suggested by the software (Van der Auwera et al. 2013).

### Identification and annotation of A-to-I editing sites

VarScan (version 2.4.3) was utilized to call variants to identify single-nucleotide variations (SNVs) (Koboldt et al. 2012). Given that inosine (I) is interpreted as guanosine (G) during reverse transcription and translation, A-to-I RNA editing is also referred to as A-to-G RNA editing (Goldstein et al. 2017; Xiang et al. 2018). Filtering was performed using a standard bioinformatic pipeline described in our previous study (Tao et al. 2021; Zhang et al. 2022). Annotation was carried out using the Ensembl Variation Effect Predictor (VEP) (McLaren et al. 2016). Briefly, only SNVs that met the following criteria were kept: base quality ≥ 25, total sequencing depth ≥ 10, alternative allele depth ≥ 2, and alternative allele frequency (AAF) ≥ 1%. SNVs that met the following criteria were filtered and removed unless annotated in the REDIportal V2.0 database (Picardi et al. 2017; Mansi et al. 2021): (1) located within the mitochondria DNA, simple repeat sequences, or homopolymer runs ≥ 5 nucleotides (nt); (2) located within six nt of splice junctions or one nt of insertions-deletions (INDELs); (3) annotated in the dbSNP database as known variant types; (4) with an AAF equal to 100% or ranging between 40% and 60% in over 90% of samples; (5) covered by high-quality reads < 10, or edited reads < 2, or editing levels < 1%.

### Identification of differential RNA editing sites

Generalized Linear Models (GLM) and Likelihood Ratio Tests (LRT) were used to compare RNA editing levels between embryonic and adult stages. Differential RNA editing (DRE) sites were identified using the following criteria: (1) differential editing levels ≥ 1%; (2) false discovery rate (FDR) < 0.05 (Srivastava et al. 2017).

### Quantification of gene expression

FeatureCounts (Version 2.0.1) was utilized to extract pseudo-counts of gene expression from BAM files (Liao et al. 2014). Subsequently, EdgeR (Version 3.7) was used to calculate transcripts per kilobase million (TPM) values for each gene, and differential gene expression analysis was performed using the Generalized Linear Model (GLM) (Robinson et al. 2010). *P*-values < 0.05 were determined to be statistically significant. The correlation coefficients (*r*) between RNA editing level and gene expression were calculated using Pearson correlation analysis.

### Enrichment analysis of gene function and pathways

Enrichr (https://maayanlab.cloud/Enrichr/) (Kuleshov et al. 2016) and DAVID online prediction tools (https://david.ncifcrf.gov/tools.jsp) (Huang da et al. 2009) were used to perform Gene Ontology (GO) biological process enrichment analysis. Items with a *P*-value < 0.05 were considered to be statistically significant.

### Prediction and mapping of RNA-binding protein binding sites

The RBPmap website (http://rbpmap.technion.ac.il) was used to predict the binding patterns of RNA-binding proteins (RBPs) to differential A-to-I RNA editing sites (Paz et al. 2014). The results of the RBPs binding prediction were visualized with the WordCloud package (Version 2.6).

### Analysis of differential alternative splicing

The replicate multivariate analysis of transcript splicing (rMATS) (Version 4.0.2) was used to detect differential alternative splicing (DAS) events across different tissues (Shen et al. 2014). The obtained alternative splicing (AS) events were classified into five categories: skipped exon (SE), retained intron (RI), alternative 5′ splice sites (A5SS), alternative 3′ splice sites (A3SS), and mutually exclusive exons (MXE). For each AS event, junction reads from inclusion and exclusion groups were averaged across replicates and used to calculate the Inclusion Level Difference (ΔPSI). DAS events were identified using the following criteria: (1) | ΔPSI | > 10%; (2) *P*-values < 0.05.

## Author contributions

**Jia-Qi Pan:** Conceptualization, Data curation, Formal analysis, Writing - original draft. **Xu-Bin Pan:** Conceptualization, Methodology, Software. **Yan-Shan Liu**: Methodology, Software, Supervision. **Yun-Yun Jin:** Methodology, Project administration, Supervision. **Jian-Huan Chen:** Conceptualization, Supervision, Funding acquisition, Writing – review & editing.

## Declarations

All authors declare that they have no known financial or personal relationships that might influence the publication of this paper.

