## Supplementary Figures for "Multi-tissue transition of A-to-I RNA editing pattern and its regulatory relevance in transcription, splicing, and translation during development"

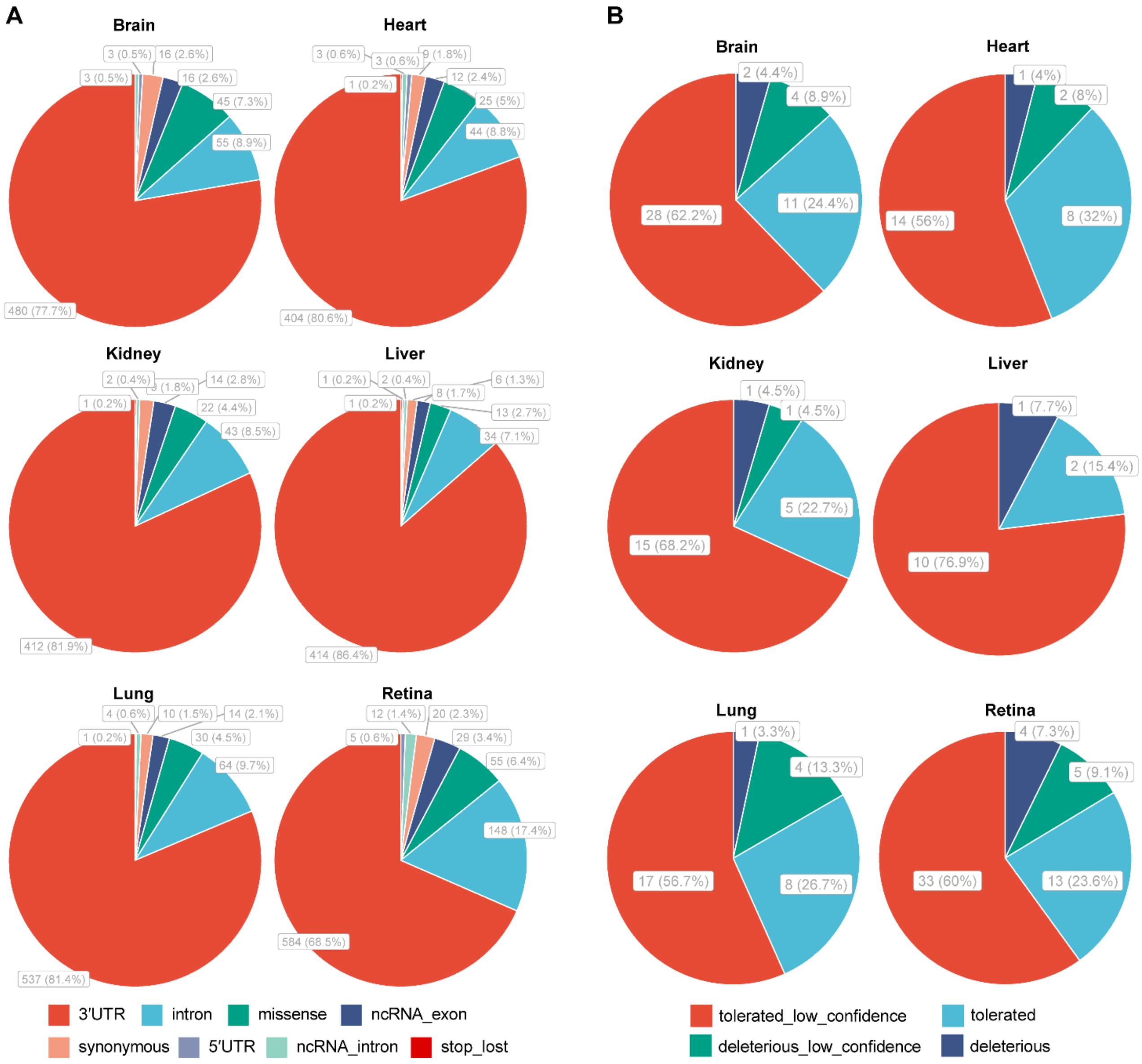


**Figure S1. Characterization of A-to-I RNA editing sites across different tissues.** **(A)** Pie charts showing the numbers and percentages of A-to-I RNA editing sites in each functional category. **(B)** Pie charts showing the numbers and percentages of missense variants in each SIFT type.

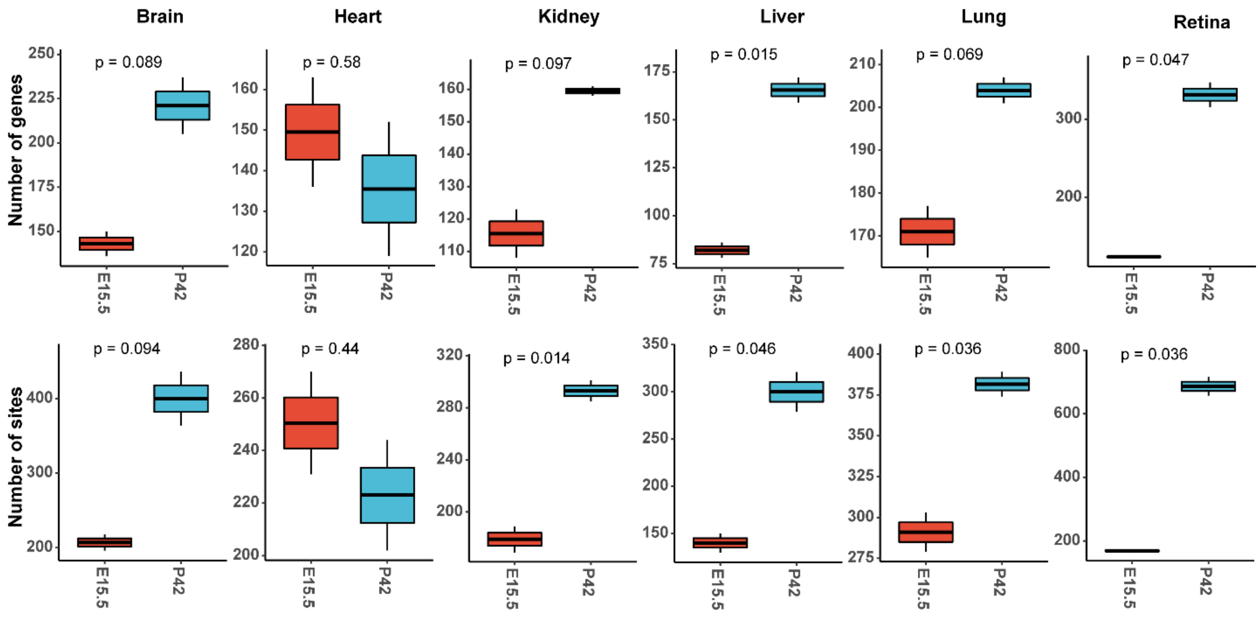


**Figure S2. Boxplot showing number of A-to-I RNA editing genes and sites between embryonic and adult stages across different tissues.**

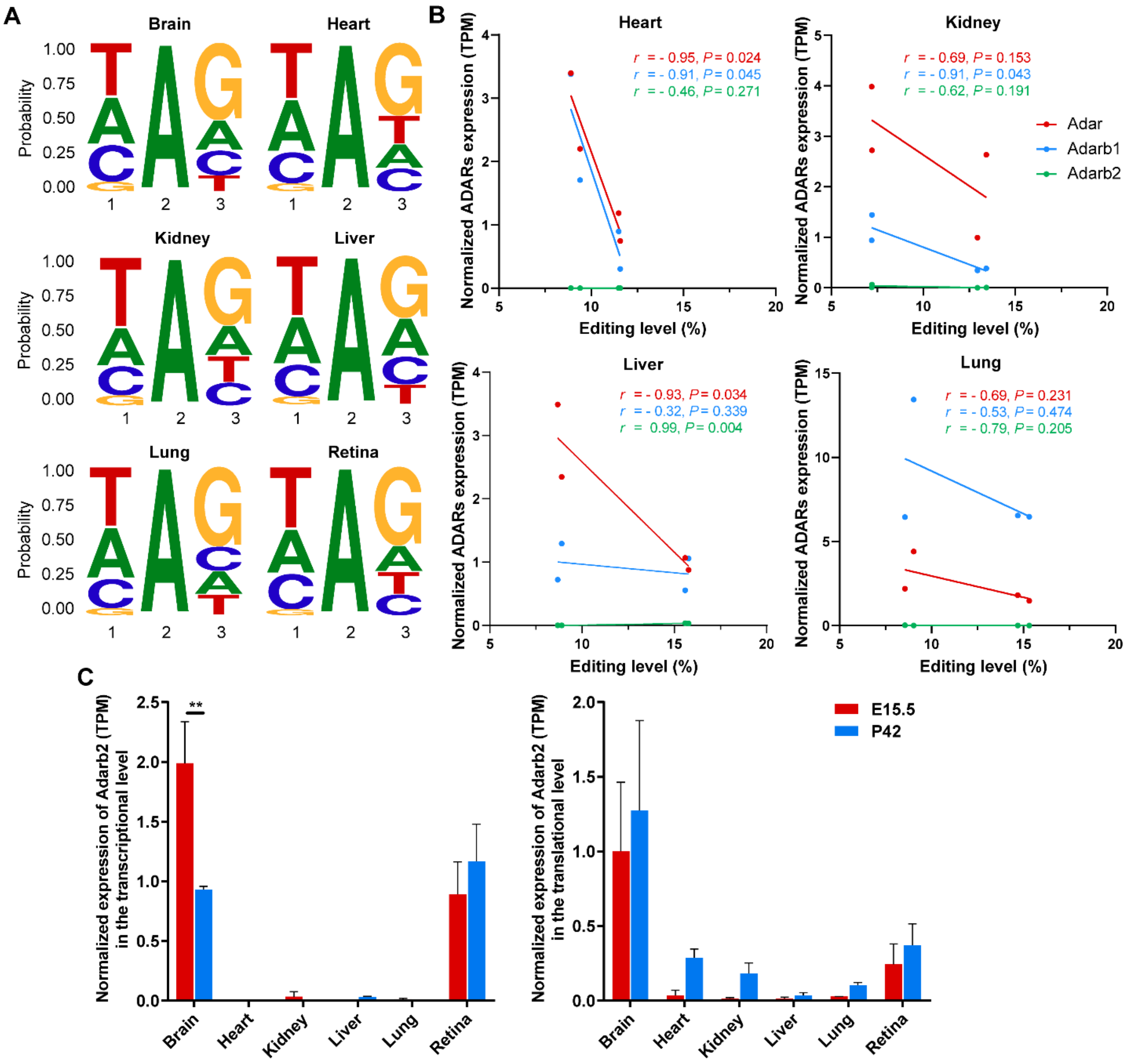


**Figure S3. (A)** Characterization of motif context around differential A-to-I RNA editing sites in different tissues. **(B)** Pearson's correlation analysis between editing levels and normalized ADARs expression in heart, kidney, liver and lung. **(C)** Normalized expression of Adarb2 in the transcriptional (left) and translational level (right).

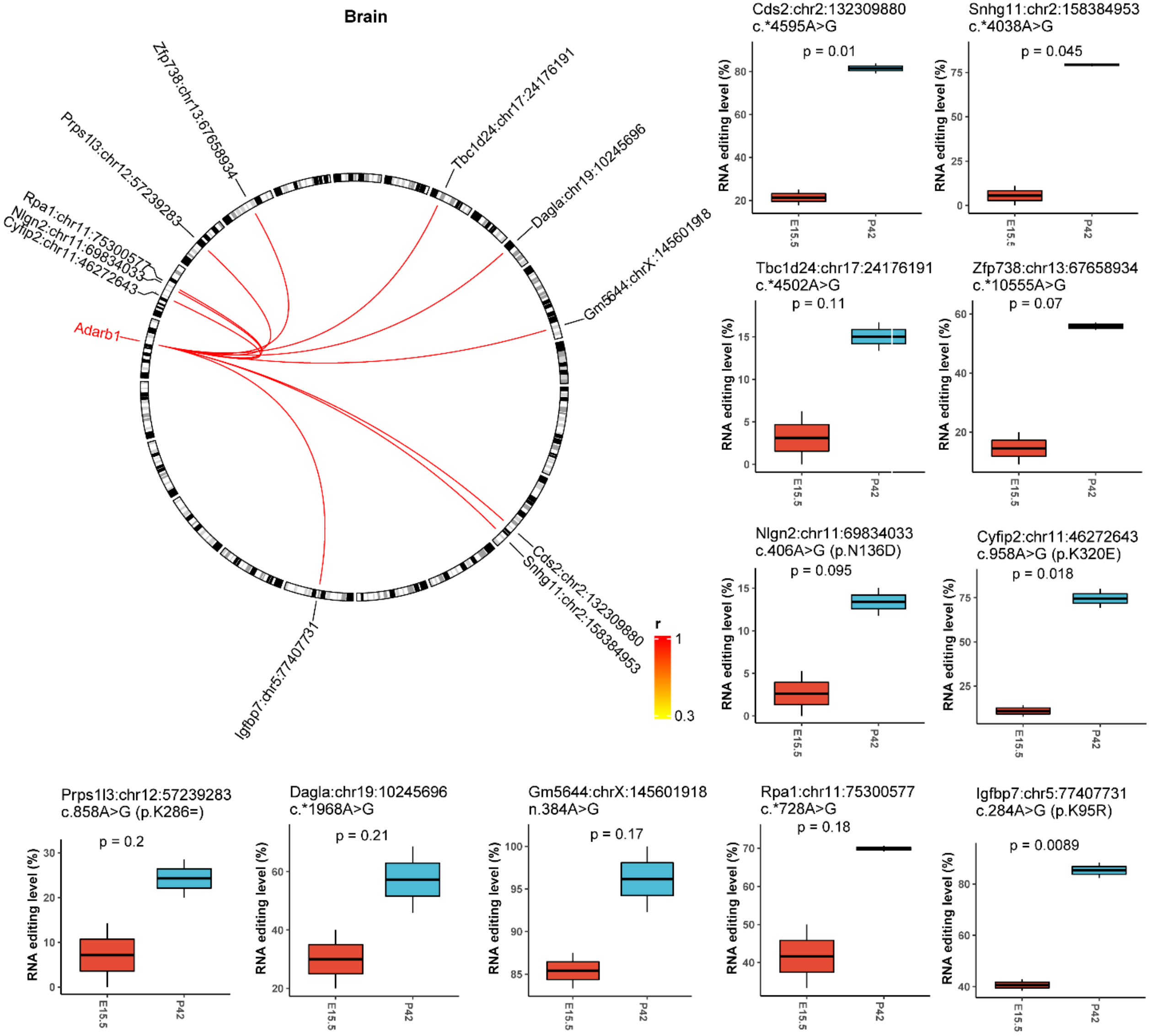


**Figure S4. The correlation between Adarb1 expression and differential A-to-I RNA editing sites in brain.**

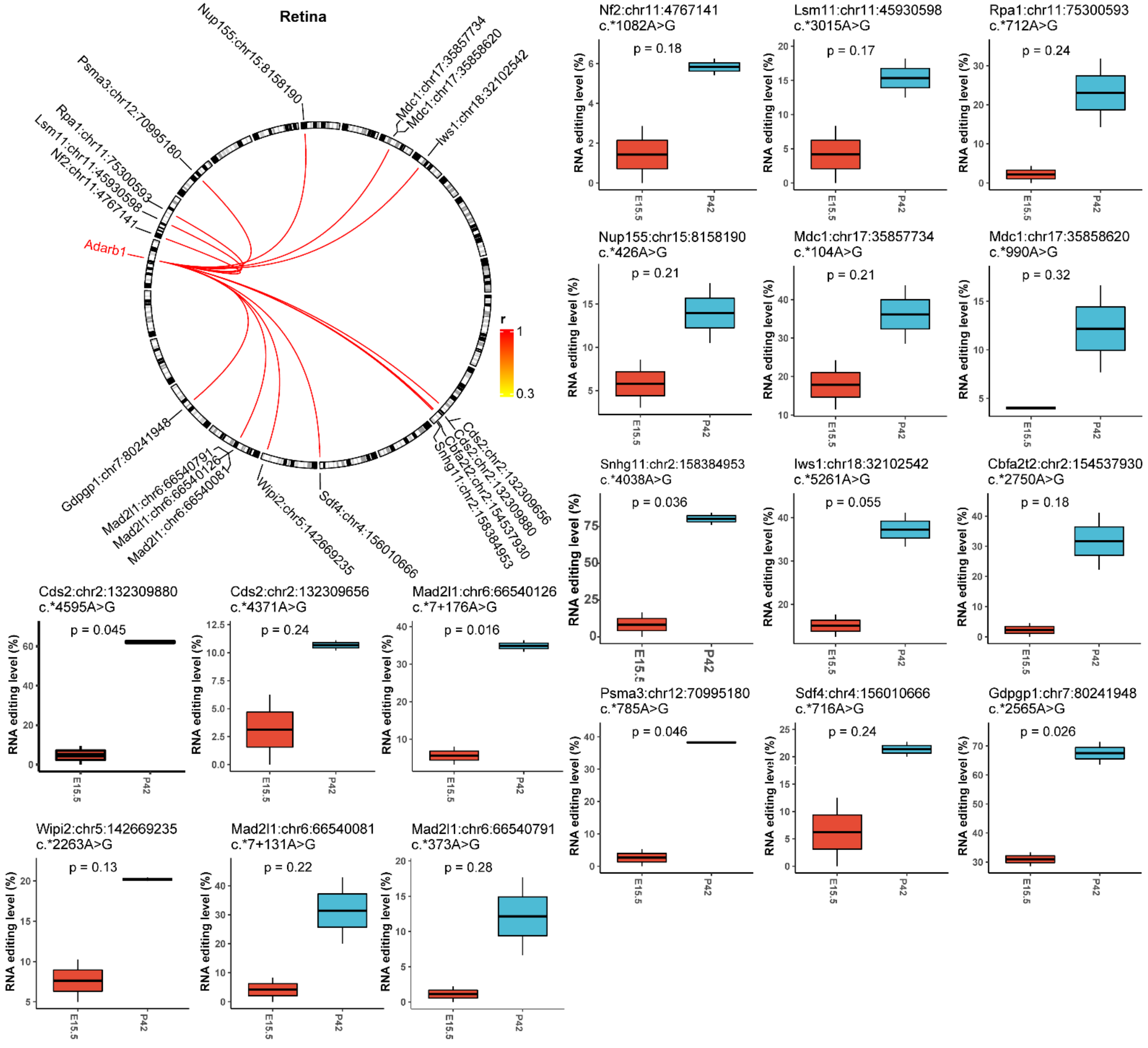


**Figure S5. The correlation between Adarb1 expression and differential A-to-I RNA editing sites in retina.**

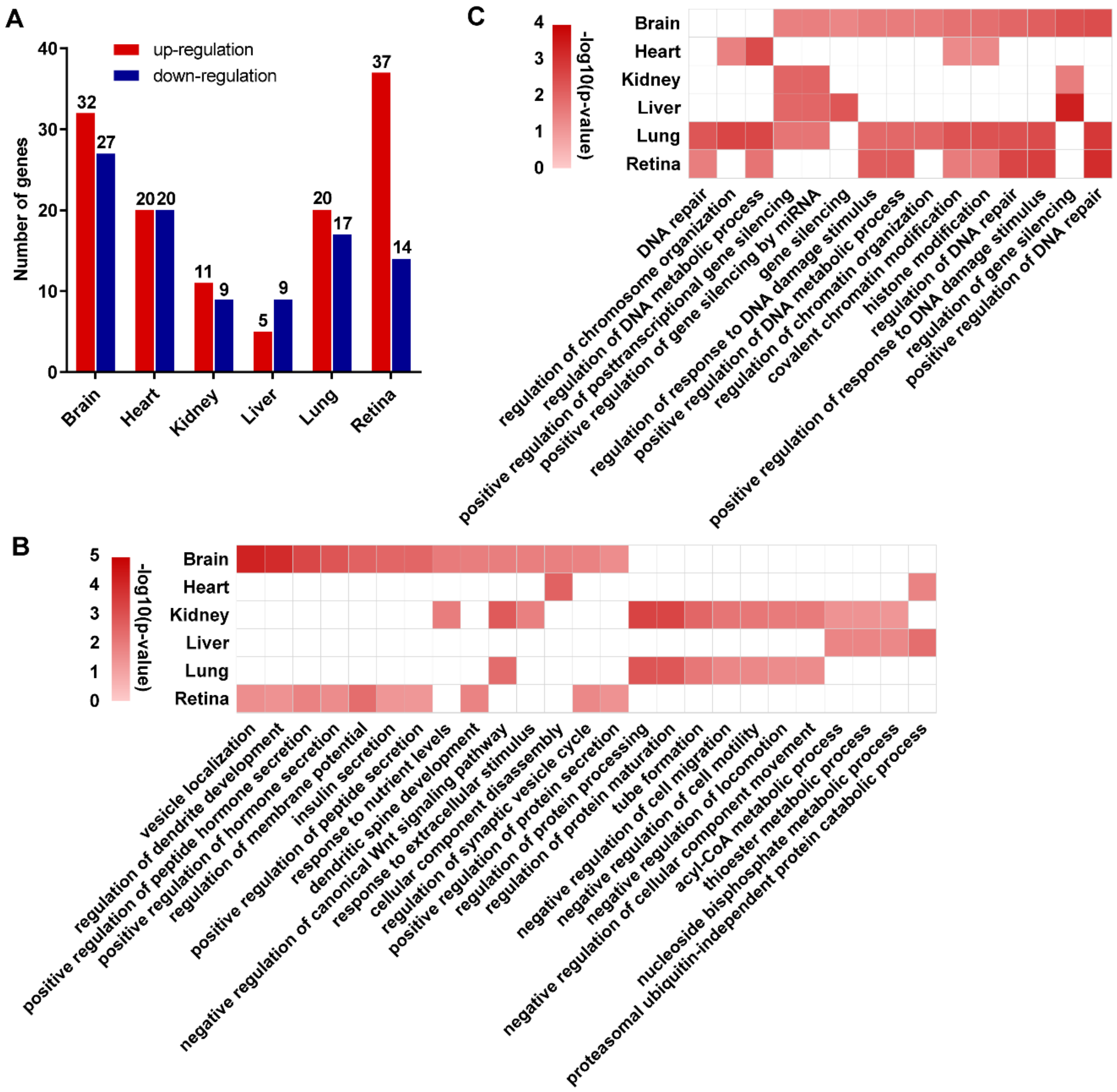


**Figure S6. DRE genes with transcriptional differential expression only. (A)** Bar plot showing the number of DRE genes with transcriptional differential expression only. Heatmap showing the enriched GO terms for DRE genes with transcriptional up-regulated **(B)** and down-regulated expression **(C)** in at least two tissue types.

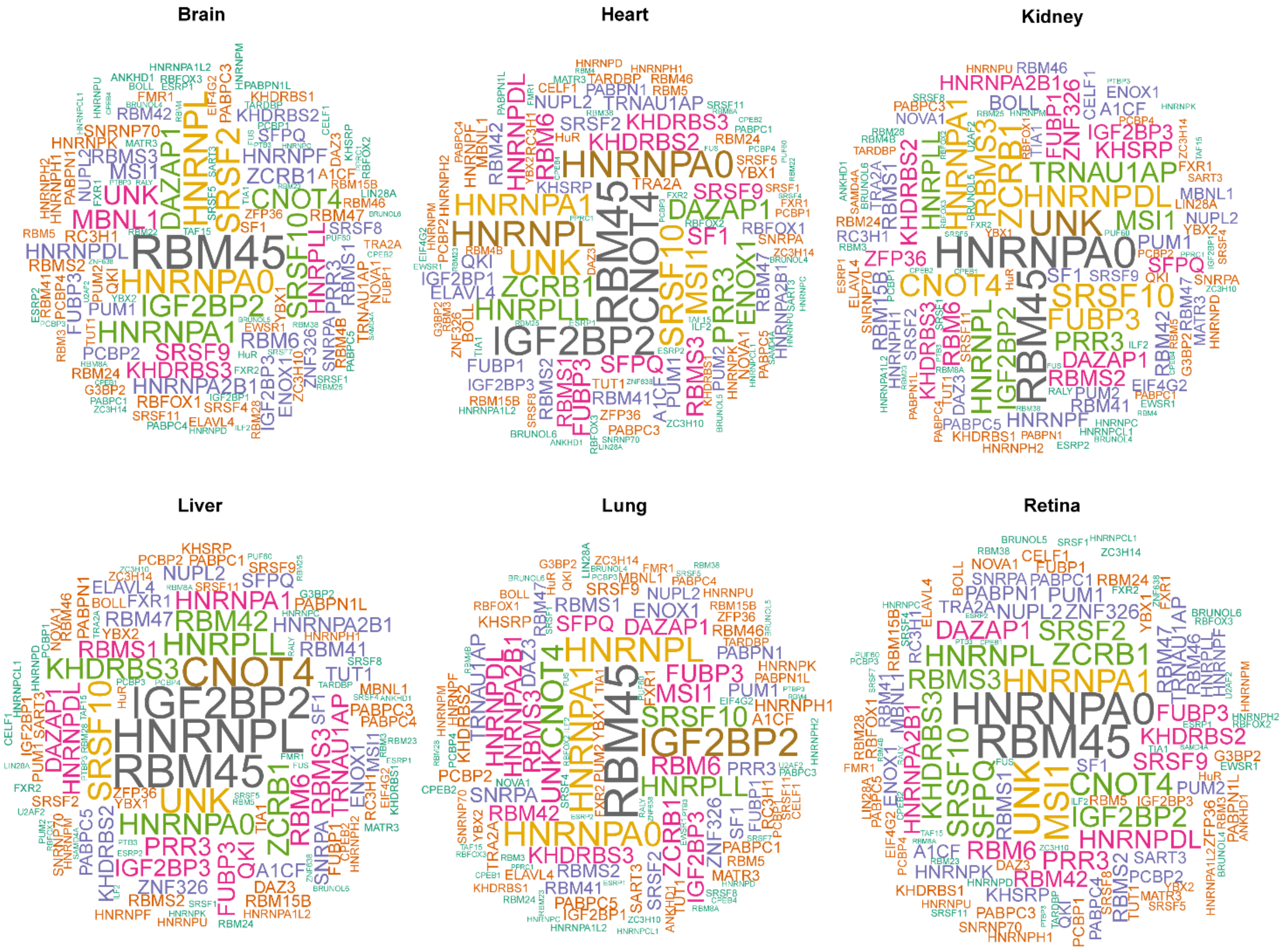


**Figure S7. RNA binding proteins (RBPs) of differential A-to-I RNA editing sites in different tissues.**

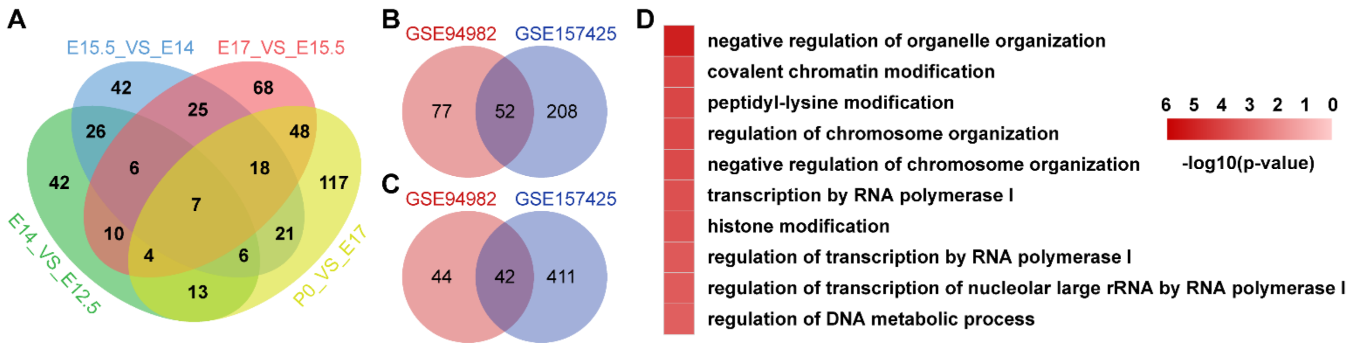


**Figure S8. Validation of RNA editing pattern during brain development in dataset GSE157425 and GSE169457. (A)** Venn plots comparing DRE sites between adjacent developmental time points. **(B)** Venn plot comparing DRE genes with differentially expressed both in mRNA-Seq and Ribo-Seq between datasets GSE94982 and GSE157425. **(C)** Venn plot comparing DRE sites with differentially expressed both in mRNA-Seq and Ribo-Seq between datasets GSE94982 and GSE157425. **(D)** Heatmap showing the enriched GO terms of DRE genes in dataset GSE157425.
